# Biosynthesis of the Antibiotic Nonribosomal Peptide Penicillin in Baker’s Yeast

**DOI:** 10.1101/075325

**Authors:** Ali R. Awan, Benjamin A. Blount, David J. Bell, Jack C. H. Ho, Robert M. McKiernan, Tom Ellis

## Abstract

Fungi are a valuable source of enzymatic diversity and therapeutic natural products including antibiotics. By taking genes from a filamentous fungus and directing their efficient expression and subcellular localisation, we here engineer the baker’s yeast *Saccharomyces cerevisiae* to produce and secrete the antibiotic penicillin, a beta-lactam nonribosomal peptide. Using synthetic biology tools combined with long-read DNA sequencing, we optimise productivity by 50-fold to produce bioactive yields that allow spent *S. cerevisiae* growth media to have antibacterial action against *Streptococcus* bacteria. This work demonstrates that *S. cerevisiae* can be engineered to perform the complex biosynthesis of multicellular fungi, opening up the possibility of using yeast to accelerate rational engineering of nonribosomal peptide antibiotics.

---

Many important therapeutics including key antibiotics are derived from compunds produced by fungal organisms^1^, yet fungal enzymatic diversity is a largely untapped resource for cheap biosynthesis of medical molecules and the potential for the discovery of novel therapeutics^2, 3^. The baker’s yeast *Saccharomyces cerevisiae* is the most well-characterised unicellular fungi and is also one of the main organisms used industrially for engineered biosynthesis^4^. Since the advent of synthetic biology, *S. cerevisiae* has been genetically reprogrammed with diverse enzymes from bacteria and eukaryotes to produce hundreds of different molecules of industrial and therapeutic relevance including opiates and anti-malarial terpenoids^5^-^9^. A major class of bioactive molecules found in fungi as well as in bacteria are the non-ribosomal peptides (Nrp). These complex molecules, produced by large assembly-line enzymes called non-ribosomal peptide synthetases (NRPS) include many front-line antibiotics including the classic beta-lactam antibiotic penicillin, naturally produced by filamentous fungi^10, 11^. However, despite a wide range of known fungal NRPS genes and the plethora advanced genetic tools for *S. cerevisiae*, there have been no reports of engineered beta-lactam antibiotic production from this yeast.

Previously, Siewers *et al.* showed that co-expression of the *Penicillium chrysogenum* NRPS gene *pcbAB* and an NPRS activator gene in *S. cerevisiae* led to cytosolic synthesis of amino-adipyl-cysteinyl-valine (ACV), the Nrp intermediate in the five gene *P. chrysogenum* pathway of penicillin G (benzylpenicillin) biosynthesis^12^. While this revealed that the first step of beta-lactam production was possible in *S. cerevisiae*, the pathway to producing an active Nrp antibiotic was incomplete. Using modular genetic tools to control enzyme expression, we demonstrate here that correct subcellular localisation, along with expression optimisation of the full five gene fungal pathway enables *S. cerevisiae* to synthesise benzylpenicillin. We further demonstrate that bioactive benzylpenicillin is secreted by engineered yeast and that spent culture media from the yeast has antibiotic activity against *Streptococcus* bacteria.

## Results

### Establishing biosynthesis and secretion of benzylpenicillin in *S. cerevisiae*

The benzylpenicillin pathway in *P. chrysogenum* consists of five genes converting cysteine, valine and the non-canonical amino acid alpha-aminoadipic acid (AAA) into a beta-lactam antibiotic via ACV (Fig 1a). The 11.3 kbp NRPS gene *pcbAB* and the NRPS activator gene *npgA* required to produce ACV were first integrated into the *S. cerevisiae* BY4741 *TRP1* locus by inserting the two genes and the bidirectional GAL1/GAL10 promoters from the previously described pESC-npgA-pcbAB plasmid^12^. ACV production in this strain (Sc.A1) was then confirmed using Liquid Chromatography Mass Spectrometry (LCMS) comparing to a chemical standard (Fig 1b). To complete the pathway and establish benzylpenicillin biosynthesis in *S. cerevisiae*, we next required efficient expression of the remaining three *P. chrysogenum* enzymes. However, as the final two steps of this pathway are known to naturally occur in the peroxisome in *P. chrysogenum* ^13^, we reasoned that cytoplasmic expression of all enzymes would not suffice and that subcellular localisation would be the key to full synthesis. We therefore took the step of tagging both *pclA* and *penDE* with the PTS1 tag^14^ to direct these enzymes to co-localise in the peroxisome upon translation. To verify expression and subcellular localisation of the enzymes, we constructed a single yeast plasmid expressing *pcbC, pclA* and *penDE* each fused to a different fluorescent protein and verified expression location by fluorescence microscopy (Fig 1c). To test whether simultaneous expression of all five genes could yield benzylpenicillin production, we removed the fluorescent protein tags from the three-gene plasmid, and transformed this into the Sc.A1 strain. The resulting strain (Sc.P1) was shown to indeed produce low quantities of benzylpenicillin when measured by LCMS (Fig 1d), and while the ACV Nrp intermediate was detected at equal amounts inside and outside cells, benzylpenicillin was found predominantly in supernatant fractions, suggesting active secretion of the antibiotic, as opposed to passive diffusion for ACV (Fig 1e).

**Figure 1.**
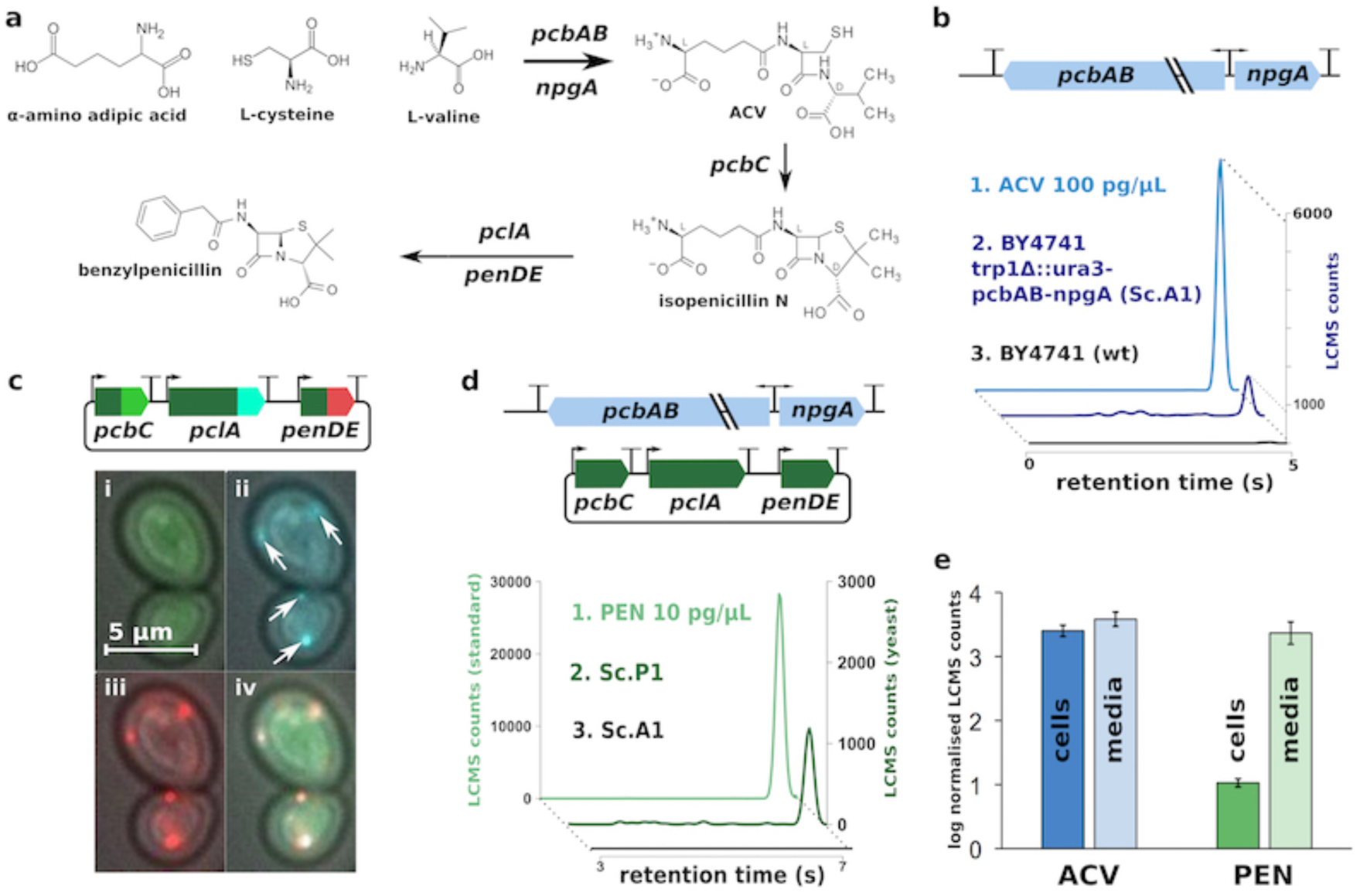
Production of benzylpenicillin by engineered *S. cerevisiae*. **(a)** Benzylpenicillin biosynthesis requires a five enzyme pathway where alpha-aminoadipic acid (AAA), cysteine and valine are converted to the tripeptide ACV by *pcbAB* and *npgA.* ACV is then converted to benzylpenicillin by *pcbC*, *pclA* and *penDE*. **(b)** Strain Sc.A1 (*S. cerevisiae* BY4741 with genomically-integrated *pcbAB* and *npgA* genes) produces ACV as observed by LCMS analysis. LCMS counts vs. retention time is shown for an ACV standard (light blue) as well as for supernatant from an Sc.A1 culture (dark blue) and a BY4741 culture (black), both supplemented with 5 mM AAA. **(c)** Subcellular localisation of enzymes encoded by *pcbC*, *pclA* and *penDE* were verified by transforming *S. cerevisiae* BY4741 with a plasmid where each gene is tagged by fusion to a different fluorescent protein. Fluorescence microscopy images show distribution for each fluorescent tag (i–iii) and a composite (iv). Arrows indicate peroxisomes. *pclA* and *penDE* genes are tagged for peroxisomal expression and *pcbC* is cytosolic. **(d)** To produce benzylpenicillin strain Sc.A1 was transformed with a plasmid expressing *pcbC*, *pclA* and *penDE* to make strain Sc.P1. LCMS counts vs retention time is shown for a benzylpenicillin standard (light green) as well as for supernatant from an Sc.P1 culture (dark green) and an Sc.A1 culture (black), both supplemented with 5 mM AAA and 0.25 mM phenylacetic acid. **(e)** Benzylpenicillin is secreted as shown by concentration-normalised log10 values of LCMS counts for cell pellets and media for both ACV and benzylpenicillin from Sc.P1 cultures grown as in (d). Error bars show standard deviation for three biological replicates.

### Pathway optimisation with combinatorial library analysis by nanopore sequencing

Next, we sought to optimise production of benzylpenicillin in *S. cerevisiae*. One common strategy for improving biosynthesis yields is to alter the expression levels of the pathway enzymes, typically by changing the promoters for the corresponding genes^15^. This can increase the flux through the pathway preventing the build-up of any inhibitory intermediates but can also aid in finding efficient expression levels of enzymes that aid their correct folding and subcellular localisation, both important considerations for this case. Therefore, using a recently described modular cloning toolkit for yeast^16^ we constructed and tested hundreds of different combinations of the benzylpenicillin pathway genes with different promoters known to vary in strength and expression dynamics (Fig 2). We used this approach to first optimise the production of the ACV Nrp intermediate, and then the conversion of ACV to benzylpenicillin. For the optimisation of ACV production, we cloned the two genes for ACV biosynthesis with different pairs of promoters on a low-copy centromeric plasmid and measured ACV yields from transformed BY4741 yeast by LCMS. All strains constructed outperformed our Sc.A1 strain (~20 ng/mL) and one combination, *pcbAB* with pTDH3 promoter and *npgA* with pPGK1 promoter, generated a strain (Sc.A2) that even outperformed yeast with the high-copy number pESC-npgA-pcbAB 2-micron plasmid containing strong galactose-inducible promoters (~280 ng/mL vs ~70 ng/mL). We attribute these higher yields to increased enzyme expression, presumably due to AT-rich sequences immediately upstream of start codons on our plasmids. These sequences are known to improve translation initiation efficiency in *S. cerevisiae*^17^.

**Figure 2.**
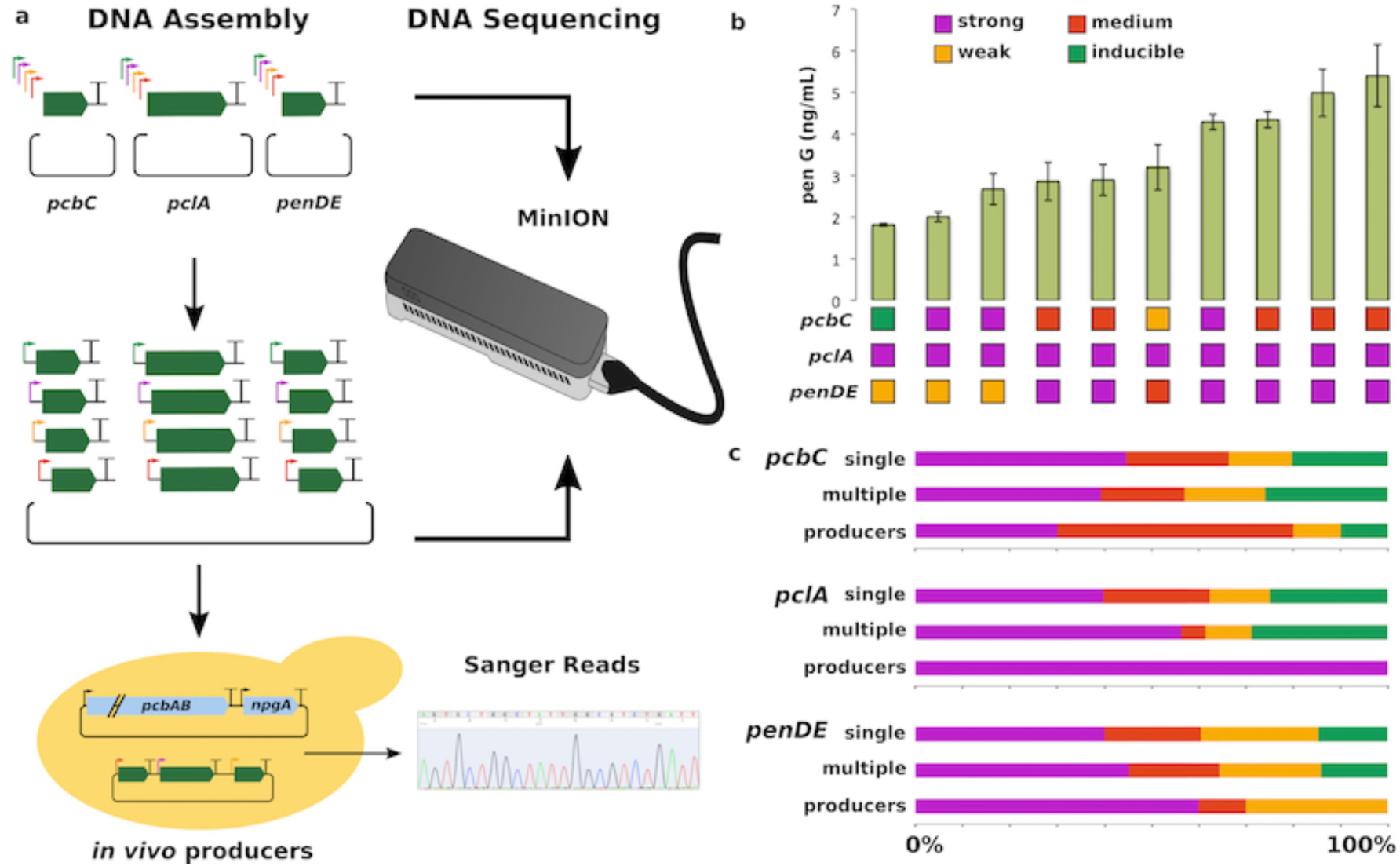
Optimisation of benzylpenicillin production in *S. cerevisiae*. **(a)** Construction and sequencing of a library with alternative promoters for *pcbC*, *pclA* and *penDE*. Ten different promoters of four categories (strong, medium, weak or inducible) were randomly used to drive expression of each of the genes responsible for conversion of ACV to benzylpenicillin. Plasmid libraries of single genes with different promoters were initially constructed and then pooled to randomly assemble a multigene plasmid library. Single gene and multigene libraries were subjected to MinION nanopore sequencing. The multigene library was transformed into ACV producer Sc.A2 and transformants producing benzylpenicillin were Sanger sequenced to reveal identity of promoters. **(b)** Concentrations of benzylpenicillin secreted into the supernatant by the ten yeast strains selected from the promoter screen. The best producing strain makes ~5 ng/mL of benzylpenicillin as determined by LCMS. Under each bar, the category of the promoter in front of each gene is shown. Error bars represent standard error of the mean from three biological replicates. **(c)** Comparison of the percentage of promoters from each of the four categories for each pathway gene during Golden Gate library construction (single and multiple gene steps) and among the ten benzylpenicillin producing yeast strains as determined by DNA sequencing.

To optimise conversion of ACV to benzylpenicillin, we next exploited one-pot combinatorial DNA assembly using Golden Gate cloning to make a diverse library of high-copy plasmids in which genes *pcbC*, *pclA* and *penDE* are each expressed from one of ten randomly assigned promoters that span a range of strengths (Fig 2a). Eight different constitutive promoters and two galactose-inducible promoters were included. The constitutive promoters were classified as “strong” (4 total), “medium” (2 total) or “weak” (2 total) promoters according to their characterisation in a previous study (Table S7)^16^. The resultant plasmid library with a theoretical diversity of 1000 members was transformed into the Sc.A2 strain and a total of 160 colonies were individually screened by LCMS for benzylpenicillin production in YPD media (Table S2).

From the screened colonies, the plasmids from ten strains showing detectable production of benzylpenicillin were sequenced to determine the promoter combinations for *pcbC*, *pclA* and *penDE* that direct efficient biosynthesis in *S. cerevisiae*. Sequencing revealed an apparent over-representation of strong constitutive promoters at the *pclA* gene (10 out of 10) and of medium constitutive promoters at the *pcbC* gene (5 out of 10) in strains with benzylpenicillin production (Fig 2b). These results suggested that either a strong level of *pclA* expression is always required for benzylpenicillin production in *S. cerevisiae*, or that our combinatorial DNA assembly had an unintended bias for incorporation of these promoter parts into the final plasmid library. To rule out any bias in our library assembly we used a MinION DNA sequencer (Oxford Nanopore Technologies) to provide long-read sequencing of the assembly products pooled from all stages in our plasmid library construction (Fig 2a). The ability of nanopore sequencing to routinely return read lengths above 5 kb make it well suited to determining the full-length products of modular DNA construction^18^. Two-directional (2D) reads of products from multigene assembly revealed *in vitro* construction of over one hundred plasmids containing different combinations of promoters in front the three pathway genes. No significant bias for incorporation was seen during either of the DNA assembly steps compared to the distribution expected by chance (Fig 2c), whereas a clear bias was seen for strong promoters being required for *pclA* in all yeast strains able to perform benzylpenicillin biosynthesis (Fig 2c).

### Antibiotic action of the spent yeast culture media

Optimisation of the biosynthesis pathway through combinatorial cloning with alternative promoters led to a >50-fold increase in yields of secreted benzylpenicillin compared to our first strain Sc.P1. The highest producer, strain Sc.P2, secretes benzylpenicillin into its growth media at a concentration calculated to be around 5 ng/mL (Fig 2b). Using this strain, we next sought to test whether benzylpenicillin secreted by engineered *S. cerevisiae* shows expected bioactivity. The secreted levels of benzylpenicillin are similar to those required for antibiotic action against various bacteria such as *Streptococcus pyogenes*^19^. Therefore, to test the bioactivity of the secreted benzylpenicillin, we diluted liquid cultures of *S. pyogenes* with spent culture media from either Sc.P2 or from an inactivated version of Sc.P2 (Sc.P2x) that does not produce benzylpenicillin (Fig 3a). Overnight growth of *S. pyogenes* was almost completely inhibited in the presence of spent culture media from Sc.P2. By contrast, spent culture media from the inactivated Sc.P2x strain resulted in normal growth of *S. pyogenes* (Fig 3b). Simultaneous exposure of *S. pyogenes* to a range of concentrations of benzylpenicillin standards revealed that the spent culture media from Sc.P2 had the antibiotic action equivalent to 3 ng/mL of benzylpenicillin being present. This concentration was then verified in parallel by LCMS analysis of the Sc.P2 culture media, demonstrating that the benzylpenicillin secreted by *S. cerevisiae* has the same bioactivity as the commercially-obtained standard.

**Figure 3.**
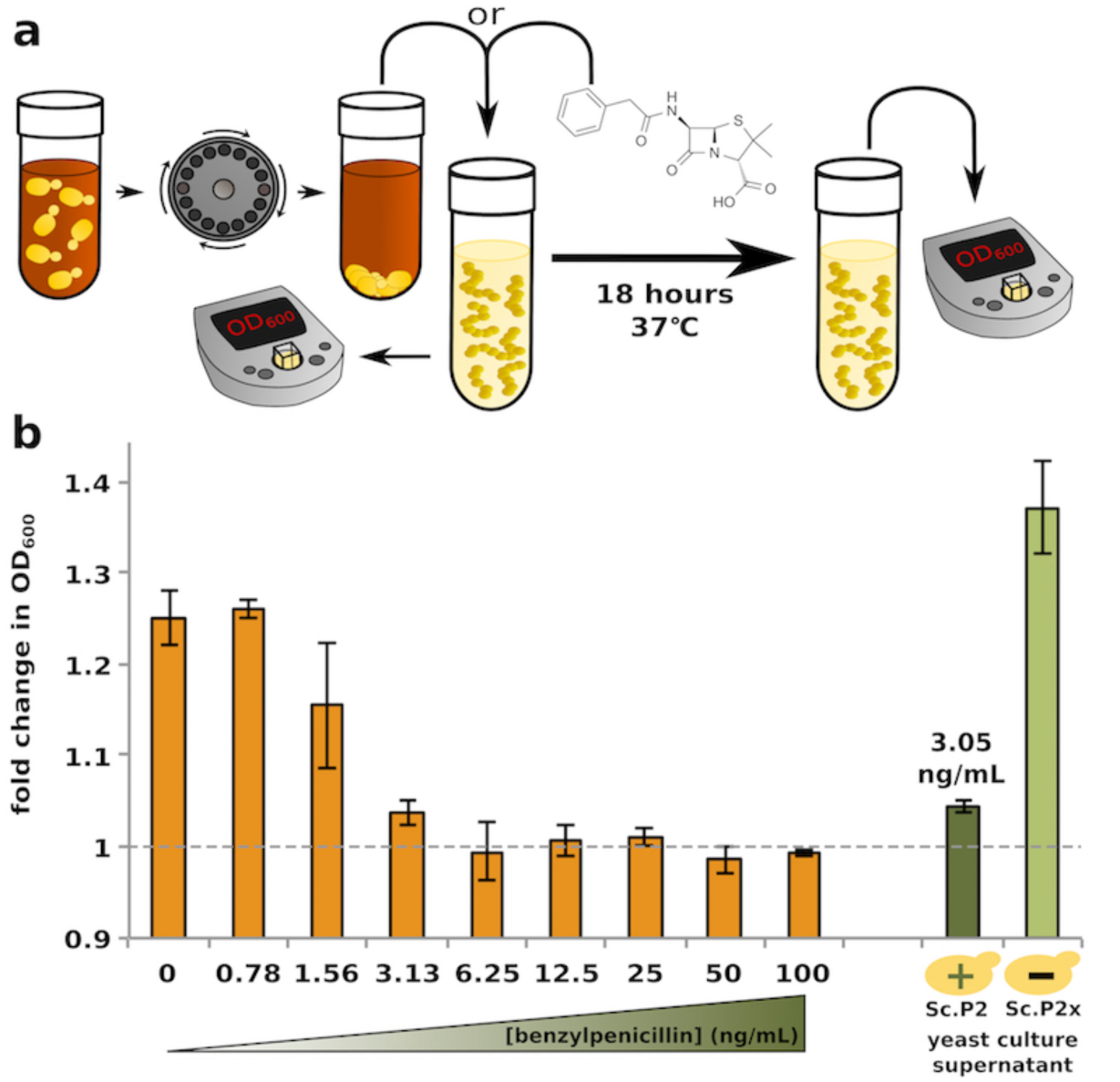
Secreted benzylpenicillin is a bioactive antibiotic. **(a)** Schematic of the spent culture media antibiotic activity test. **(b)** Fold change in OD600 of *S. pyogenes* cell cultures grown either in the presence of increasing concentrations of benzylpenicillin standard over 18 hours (orange bars, left) or in the presence of supernatant from benzylpenicillin producing yeast Sc.P2 (dark green bar, plus sign) and yeast Sc.P2x with an inactive *pcbAB* gene (light green bar, minus sign). Concentration of benzylpenicillin in Sc.P2 supernatant as measured by LCMS is given above the dark green bar. Error bars show standard error of the mean.

## Discussion

Our results demonstrate the first biosynthesis of a bioactive nonribosomal peptide (Nrp) antibiotic from engineered *S. cerevisiae* yeast. After achieving production of benzylpenicillin, we improved yields by optimising pathway enzyme expression and exploited long-read nanopore sequencing to verify the combinatorial DNA assembly of our libraries to ascertain the best combination of promoter strengths for increased yields. We observed that this always required strong expression of the peroxisomally-located enzyme *pclA.* As engineered heterologous biosynthesis of penicillin has only previously been observed with *Hansenula polymorpha*, a methylotrophic yeast with large peroxisomes^20^, this suggests that boosting peroxisomal expression further, for example by increasing peroxisome formation, by may be a future approach to improve yields from *S. cerevisiae.* Further yield improvements could also be achieved using other metabolic engineering strategies^21, 22^.

More generally, we view this achievement in *S. cerevisiae* as a paradigm for establishing complex fungal biosynthesis pathways in a standard chassis organism used extensively in research and industry. Give the impressive wealth of genetic tools and the ease of engineering with this microbe, we see it as the ideal testbed for screening and diversifying fungal pathway biosynthesis. We especially anticipate that expression and genetic manipulation of other bioactive Nrp pathways in *S. cerevisiae* could advance the screening of engineered or newly-discovered nonribosomal peptide synthetase (NRPS) enzymes from fungi. NRPS enzymes represent a particularly attractive class of enzymes for engineering efforts as they are inherently modular with each module region recognising one amino acid and incorporating it into the Nrp product^15, 23^. NRPS modules incorporate both the standard 20 amino acids, as well as hundreds of non-proteinogenic amino acids, including D-enantiomers^24^. Thus by combining different modules together it should be possible make chimeric NRPS enzymes that produce thousands of novel Nrp molecules predisposed by evolution for bioactivity^25^. However, determining the structural boundaries of modules within natural NRPS enzymes has so far proved extremely challenging due to the difficulty in obtaining structures for these massive enzymes^23, 26^. An alternative approach to structure-guided mutagenesis is exhaustive mutation and screening. Simple biological screens such as those based on the antibiotic activity shown in this study could help pave the way towards true combinatorial biosynthesis of novel bioactive Nrp molecules. Such endeavors are extremely important in light of widespread resistance to antibiotics and the decreasing number of new antibiotics developed through traditional means^27^.

## Acknowledgements

The authors are grateful to Dr Verena Siewers and Prof Jens Nielsen for their donation of the pESC-npgA-pcbAB plasmid, and would like to thank Prof Shiranee Sriskandan and Kristin Krohn Huse for their help handling *S. pyogenes* for the growth inhibition assay.

## Funding

This work was funded in the UK by BBSRC awards BB/K006290/1 and BB/K019791/1 and EPSRC award EP/L011573/1

## Methods

### Construction of strains

Strain Sc.A1 was made by replacing the TRP1 gene in BY4741 with the *pcbAB*-*npgA* segment from the pESC-*npgA*-*pcbAB* plasmid from a previous study^12^. A URA3 gene from *K. lactis* was integrated upstream of the *pcbAB* gene to allow genomic integration and the use of media without uracil was used to enable comparison of the ACV production of this strain with that of a BY4741 strain transformed with the ura-marked pESC-*npgA*-*pcbAB* plasmid. A full genetic map of the altered TRP1 locus is provided as an annotated Genbank file in the Supplementary Sequences. All other strains used in all experiments were constructed by transforming plasmids into BY4741. A Zip file containing annotated Genbank files of all plasmids can be requested.

### Growth of Strains for ACV and penicillin production

For all ACV and penicillin producing experiments, cells were first grown overnight in synthetic complete media minus the appropriate amino acids for selective pressure with glucose as the carbon source. For plate based assays, cells were grown overnight at 700rpm at 30°C. For 50 mL falcon tube based assays, cells were grown at 225 rpm at 30°C. Overnight cultures were then back-diluted into production media and grown at 20°C (216 rpm) for 20 hours (for plate based assays) or until the OD600 reached between 0.6 and 0.8 (for 50 mL falcon tube assays). Table S5 details the composition of production media for different experiments.

### Fluorescence microscopy

Microscopy for Fig 1 was carried out with a Nikon Eclipse Ti, using the NIS Elements AR software. The objective was set at 60x. Slides were fixed with yeast cells to visualise. The excitation wavelengths for detection of Venus, mRuby2 and mTurquoise2 fluorescence were 535, 590 and 535 nM respectively.

### Preparation of standards and samples for LCMS

For all LCMS experiments, standards were prepared as follows. ACV standards were prepared by dissolving ACV (BACHEM H-4204) in water to a concentration of 10 ng/μL and making three 10-fold dilutions. This gave four standards with concentrations of 10 ng/μL; 1 ng/μL; 100 pg/μL; 10 pg/μL. Benzylpenicillin standards were prepared by dissolving the sodium salt of penicillin G (Sigma P3032) in water. The same concentrations were used for benzylpenicillin as were used for ACV.

Cellular extracts for LCMS for the data in Fig 1 were prepared as follows. 30 mL of cell culture was collected at an OD600 of 0.6. Cell culture was centrifuged at 7000 × g for 10 minutes, and supernatant was either kept for LCMS (as in Fig 1d) or discarded. The cell pellet was resuspended in 100 μL methanol. 50 μL of the resuspension was transferred to a microcentrifuge tube with 25 μL of glass beads (Sigma G8772-100G) on ice. The tube was then vortexed for 30 seconds and then placed on ice for 30 seconds, and these two steps were repeated three times (for a total of four sets of vortexing and incubation on ice). The tube was then centrifuged at 12,000 × g for 30 minutes, and 40 μL of supernatant was aliquoted to a separate tube for LCMS measurement. Supernatants for LCMS from cultures grown in 96 well plates in Figure 2 were obtained by centrifuging plates at 3000 × g for 30 minutes.

### LCMS

An LC/MS/MS method was developed for the measurement of penicillin-G and related materials, using an Agilent 1290 LC and 6550 quadrupole time-of-flight (Q-ToF) mass spectrometer with electrospray ionization (Santa Clara, CA). The LC column used was an Agilent Zorbax Extend C-18, 2.1 × 50mm and 1.8um particle size. The LC buffers were 0.1% formic acid in water and 0.1% formic acid in acetonitrile (v/v).

The gradient elution method is detailed in Table S6. Quantitation was based on the LC retention times of standards and the area of accurately measured diagnostic fragment ion for each molecule (Table S6). The protonated molecules of each analyte, [M+H]+, were targeted and subjected to collision induced dissociation (collision energy 16eV), with product ions accumulated throughout the analysis. Solutions of penicillin-G and ACV standards in water were used to generate calibration curves.

The linear range of the method was determined by injecting standards over a range of concentrations. The lower limit of detection (llod) was determined by the amount a sample resulting in a peak with a signal-to-noise of 3:1. The lower limit of quantitation (lloq) was taken to be the concentration of analyte that produced a signal-to-noise of 10:1. The llod and lloq for penicillin-G were found to be on-column injections of 5 pg and 20 pg respectively.

### Calculation of yield of ACV and benzylpenicillin from LCMS data

For both ACV and benzylpenicillin, pure chemical standards were run at the following concentrations: 10 ng/mL, 100 ng/mL, 10 μg/mL, 100 μg/mL. The corresponding LCMS counts for these standards were plotted against the concentrations of the standards and the linear range of the resulting plot was used to construct a line of best fit in excel. The corresponding line equation was used to obtain values for the yield in ng/mL of ACV and benzylpenicillin from experimental samples based on the LCMS counts for these molecules.

### Promoter screen for optimising conversion of ACV to penicillin

The assembly of multigene (*pcbC*, *pclA*, *penDE*) plasmids with ten randomised promoters was split into two stages: assembly of single-gene constructs, then assembly of multigene constructs. For single-gene construct assembly, an equimolar mix of all ten promoters was made with a final concentration of 50 fM (referred to as “promoter mix”). This was used as a type 2 plasmid according to the yeast toolkit specification16. Then, Golden Gate reactions were set up with the following parts according to the Yeast Toolkit cassette plasmid Golden Gate assembly protocol.

Each of the three reactions were transformed into *E. coli*, and for each of the three transformation plates, transformant colonies were mixed together into a single overnight culture each. From each of the three resulting cultures a plasmid library was prepared, of which an aliquot was used to construct a pooled sample for nanopore sequencing (see section below). The three resulting single-gene plasmid libraries were used to set up a single multigene golden gate reaction according to the yeast toolkit protocol.

This Golden Gate reaction was transformed into *E. coli*, and all transformant colonies were mixed together into a single overnight culture. A multigene plasmid library was prepared from this overnight culture. Part of this plasmid library was prepared for nanopore sequencing (see section below), while 4 μg was used to transform into *S. cerevisiae* strain Sc.A2. The resulting transformants were screened by LCMS for the production of benzylpenicillin and the promoter regions of the multigene plasmids from producer strains were identified by Sanger sequencing.

### Nanopore sequencing library construction

To enrich for the penicillin pathway assembly DNA and remove assembly vector backbone DNA, the multigene assembly library was digested with EcoRI and AlwNI. Restriction digest products ranging from 5616 to 6117 bp were isolated by agarose gel electrophoresis and purified using a QIAquick gel extraction kit (Qiagen). The single pathway gene assembly libraries were similarly enriched by digestion with BsmBI and AlwnI. Restriction digest products ranging from 2062 to 2229, 2549 to 2716 and 1885 to 20152 bp for the *pcbC*, *pclA* and *penDE* assemblies, respectively, were purified.

Enriched assembly DNA for the multigene and single gene assemblies was quantified on a Qubit 2.0 fluorometer (Thermo Fisher Scientific) using a Qubit dsDNA HS Assay Kit (Thermo Fisher Scientific). The four samples were combined to a give an equimolar mix (assuming a molecular weight for each assembly based on the mean promoter length) with a total DNA content of 2.6 µg in 45 μL dH_2_O.

DNA underwent end repair using NEBNext FFPE DNA Repair Mix (M6630, New England Biolabs) according to the manufacturer’s instructions. The repaired DNA was recovered using Agincourt AMPure XP beads (A63880, Beckman Coulter), washed twice in 200 μL 70% ethanol and eluted in 46 μL dH_2_O. DNA was then dA-tailed using NEBNext Ultra II End Repair/dA-Tailing Module (E7546, New England Biolabs) according to the manufacturer’s instructions and recovered using Agincourt AMPure XP beads as before, eluting in 30 μL dH_2_O.

The dA-tailed library DNA was then processed using Blunt/TA Ligase Master Mix (M0367, New England Biolabs) and a Nanopore Sequencing Kit (SQK-NSK007, Oxford Nanopore Technologies) according the manufacturer’s instructions to ligate adaptors and tethers to the library. 50 μL Dynabeads MyOne Streptavidin C1 beads were washed twice in buffer BBB (Nanopore Sequencing Kit) and then resuspended in 100 μL BBB. These beads were then added to the processed DNA sample and mixed for 5 minutes at room temperature. Beads were washed twice with 150 μL BBB before eluting the sample in 25 μL ELB (Nanopore Sequencing Kit). The library was quantified by Qubit as before and yielded a total of 253 ng.

### Nanopore sequencing

A fresh MinION R7 Flow Cell Mk I (FLO-MIN104, Oxford Nanopore Technologies) was loaded into a MinION MK I (MIN-MAP002, Oxford Nanopore Technologies) and primed using a Nanopore Sequencing Kit according to the manufacturer’s instructions. The sequencing mix was generated by combining 75 μL RNB, 65 μL NFW and 4 μL Fuel Mix (Nanopore sequencing kit) before adding 6 μL of the processed DNA library. This sequencing mix was loaded into the flow cell and sequenced using the 48 hour sequencing script on MinKNOW (Oxford Nanopore Technologies). After 18 hours, the script was stopped and a fresh sequencing mix was prepared and loaded into the flow cell. The 48 hour sequencing script was then restarted. This reloading process was repeated after a further 4.5 hours. The sequencing script was stopped once read acquisition had slowed to less than 1 successful read in a 5 minute period.

### Analysis of nanopore sequencing data

Oxford Nanopore’s cloud-based Metrichor application ‘2D Basecalling for SQK-MAP006 v1.69’ was used to basecall data from a MinION run that used R7.3 chemistry. Poretools28 (https://github.com/arq5x/poretools) was used to extract sequence files and the program lastal v658 (http://last.cbrc.jp/) was used to align the 2D reads to a database of all the potential promoters and all combined CDS+terminators using options -s2 -T0 -Q0 -a1 -fTAB -e50*. To remove reads corresponding to other DNA sequences present, the database was also populated with sequences for AmpRTerm_AmpR_AmpRProm (part of the single assembly plasmid backbone), His3Prom_His3Term_2Micron_KanRTerm_KanR_KanRProm (part of the multiple assembly plasmid backbone), ColE1 (part of both the single and multiple assembly plasmid backbone), and the lambda phage whole genome.

In order to build up a picture of which part of a plasmid each read represented, a custom-built script ordered alignments first by read, then by read coordinates. It then identified reads as originating from a particular plasmid by looking at plasmid regions in the database that each read had aligned to. These identified reads were required to be a similar length (within 15%) to the sum of lengths of regions of plasmids that they were aligned to. The script identified digested multiple gene assemblies, digested single gene assemblies and, due to reduced function of BsmBI, non-digested single assemblies. To be identified as a digested single gene assembly, the read was also required to start and finish within 50 bp of the start and end of the first and final regions respectively. This minimised the chance of misidentifying a multigene assembly as a single gene assembly. The promoters at each position within these identified reads were recorded.

### *S. pyogenes* growth inhibition assay

An overnight culture of *S. pyogenes* (H584, M1 type) was grown for 24 hours at 37°C (5% CO2) in Todd Hewitt Broth (THB). This culture was diluted in 10X THB to an optical density of OD600 0.2. In separate wells of an optically-transparent 96 well plate (VWR 3596) 10 μL of this solution was added to 90 μL of either a known concentration of benzylpenicillin dissolved in 0.25 mM Phenylacetic acid or 90 μL of spent culture media from Sc.P2 or Sc.P2x strains grown in production conditions. Sc.P2x is a variant of Sc.P2 known to not produce benzylpenicillin due an inactive *pcbAB* caused by mutation in the coding region of the gene’s terminal module region.

## Footnotes

* [-s2 use both query strands. -T0 local alignment. -Q0 use fasta as input. -a1 gap existence cost of 1. -fTAB tabular output -e50 minimum alignment score of 50.]

